# Whole-organism spatial transcriptomics at single-cell resolution in *C. elegans*

**DOI:** 10.64898/2026.04.09.717568

**Authors:** Jose David Aguirre, Xuan Wan, Carsten H. Tischbirek, Nicholas Markarian, Core Francisco Park, Long Cai, Paul W. Sternberg

## Abstract

Spatial transcriptomics has advanced our understanding of tissue organization by mapping gene expression in its native context yet applying these techniques to whole organisms remains a significant challenge. *Caenorhabditis elegans* is well-suited to whole organism-level analysis because its compact size, transparency, reproducible anatomy, and genetic tractability make it possible to link molecular and cellular changes to circuit function and behavior within the same animal. However, current transcriptomic approaches in *C. elegans* are often limited by spatial resolution or multiplexing capacity, making it challenging to profile multiple gene expression patterns across intact worms while preserving spatial context. Here, we present a single-molecule fluorescence *in situ* hybridization workflow that enables multiplex imaging with single-cell resolution across the entire worm. This approach allows sequential imaging of one gene per fluorescent channel using two channels across 20 hybridization rounds, enabling the profiling of up to 40 genes while preserving spatial context. We further provide a curated marker-gene panel for reproducible neuron identification, which, together with probabilistic assignment of transcripts to segmented nuclei, enables quantitative measurements of gene expression levels. We used this method to identify up to 86 neuronal classes and reveal sex- and neuron-specific expression patterns at single-cell resolution. Together, these results establish a scalable framework for the spatial analysis of gene expression and cell identity in intact *C. elegans*.

## Introduction

*C. elegans* is a versatile and well-established model system with a compact anatomy and stereotypical cell lineage. It benefits from a wealth of community resources such as a neuronal connectome, annotated cell identities, and large genomic and transcriptomic studies and databases^1–5^ describing gene expression in whole worms and at single-cell resolution. Its compact body allows the whole organism to be imaged in a single volume, while its stereotyped cell lineage allows spatial transcript patterns to be mapped to known cell identities with defined functions, a level of cellular resolution difficult to achieve in other organisms. Together, these features make *C. elegans* an attractive system for spatial transcriptomics at the level of the intact organism.

Building on the rich repertoire of existing methods for imaging gene expression in *C. elegans*, we present a robust spatial transcriptomic pipeline to image targeted gene panels to map gene expression with single-cell resolution in anatomically intact whole-mount worms. Pioneering work with smFISH in *C. elegans*^6–8^ showed individual gene expression in whole worms, and optimized protocols showed high signal-to-noise gene expression for adults as well as embryos^9–11^. More recent approaches include expansion microscopy with high-resolution segmentation of the worm morphology using light microscopy methods^12^ as well as digital reconstructions of gene expression based on sequencing information, taking advantage of the worm’s stereotypical morphology preserved on a single-cell level^13,14^.

The pipeline we introduce here uses two fluorescent channels, allowing sequential imaging of up to 20 genes per channel. This enables spatial analysis of 25 target genes together with a marker gene panel to identify cell types, without the extensive image alignment and postprocessing required for more advanced high-plex barcode-based workflows^15–21^. Our analysis pipeline circumvents the use of additional segmentation markers beyond nuclear DAPI dye staining to assign gene expression patterns to individual cell types.

We demonstrate the use of our experimental and analysis pipeline by profiling gene expression in both heads and tails of whole hermaphroditic and male worms and screening for sexually dimorphic gene expression. This toolkit is potentially suitable for a wide range of experiments, particularly for smaller gene panels where manual solution exchanges are still practical for multiplex imaging. Thus, we see several potential application cases in which a robust, multiplex imaging approach of targeted gene panels can contribute to accelerating experimental studies for *C. elegans* research, such as the generation of genetic driver lines specific to individual neuron classes.

## Results

### *C. elegans* sequential hybridization workflow

We developed a sequential smFISH workflow for multiplex detection of mRNA transcripts in anatomically intact *C. elegans*. Because the multi-layer worm cuticle is impermeable to many small molecules^22^ and limits efficient probe entry, a major part of our protocol was to increase *C. elegans* cuticle permeability. For this, we treated fixed animals with TCEP to chemically reduce the collagen fibers within the cuticle (**Fig. 1a**). TCEP is more stable and less prone to oxidation than thiol-based reducing agents like β-mercaptoethanol or DTT, and remains active across a broad pH range^23–25^. We further permeabilized the cuticle by treating the worms with SDS and by including an additional enzymatic digestion step with collagenase IV^12^.

**Figure 1.**
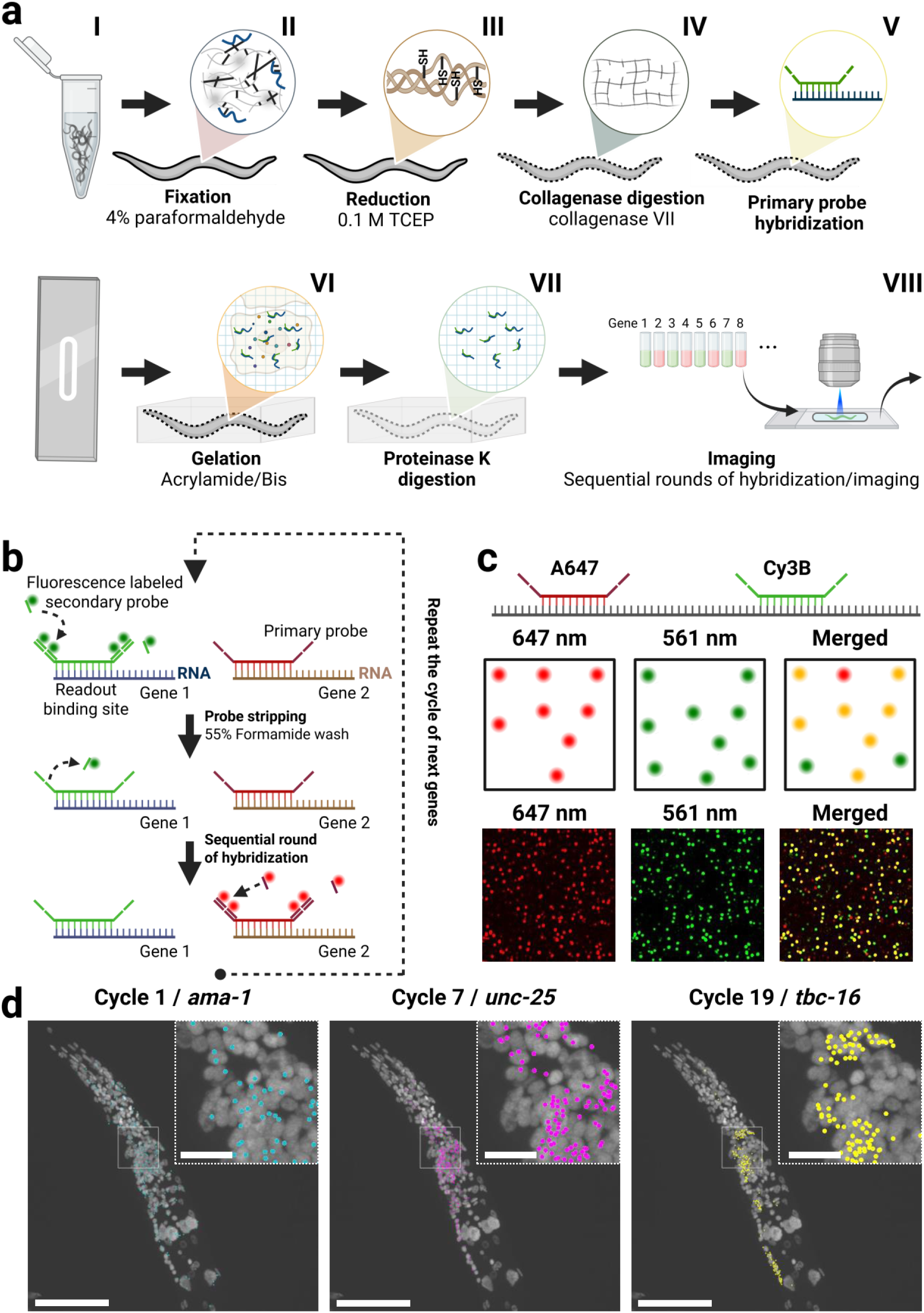
Workflow for the sequential smFISH protocol in C. elegans. (a) (I) Live animals are collected and fixed with 4% paraformaldehyde. (II) Animals are incubated in TCEP/borate buffer to reduce disulfide bonds in cuticle collagen fibers. (III) Collagenase IV treatment permeabilizes the cuticle, allowing penetration of reagents and probes. (IV) Animals are incubated with the probe oligo pool and an 18-nt poly-T acrydite-functionalized LNA oligonucleotide in hybridization buffer. (V) Hydrogel polymerization is performed on a functionalized coverslip using an activated monomer solution. (VI) Proteinase K treatment clears cellular proteins and reduces background fluorescence. (VII) Sequential rounds of readout probe hybridization and imaging are performed. (b) Schematic of sequential hybridization using primary and readout probes. Each gene is labeled with a primary probe with a unique overhang sequence complementary to a fluorescent readout probe. Readout probes are stripped with a formamide wash before the next set is hybridized and imaged. (c) A control gene (*ama-1*) was targeted with two sets of primary probes, each detected by a distinct fluorescent readout. Colocalization was assessed by comparing dots between channel 1 and 2, with dots within a 2-pixel radius considered colocalized. Scale bar, 20 μm. (d) Transcript detection across multiple hybridization cycles. Scale bar, 50 μm; inset, 10 μm.

As increased tissue permeabilization can lead to a loss of tissue integrity, we performed a gentle post-fixation step using BS(PEG)5 for tissue stabilization^26^. Samples were treated with N-(propionyloxy)succinimide to reduce non-specific binding of probes to cellular components, improving the signal-to-noise ratio for RNA detection. Primary probes were hybridized against target mRNAs, which contain two repeats of a unique overhang sequence that could be bound by complementary fluorescent probes^15–17^ (**Fig. 1b**). Acrydite-modified poly-T probes were included to anchor polyadenylated mRNAs to the hydrogel solution during polymerization^12^. To reduce the strong autofluorescence and light scattering in *C. elegans*^27–30^, the hydrogel-embedded samples were treated with Proteinase K to digest cellular proteins and lower background noise, further enhancing the signal-to-noise ratio.

During each sequential imaging cycle, we targeted a small subset of genes using fluorescently labeled “readout probes” that hybridize to the corresponding overhang sequences of their respective primary probes. Once the first round of readout probes was imaged, the readout probes were removed via a formamide wash, followed by the next imaging cycle, enabling sequential detection of different genes across multiple cycles (**Fig. 1b–d**).

### Probe specificity validation by colocalization

To validate the specificity of our primary probes, we implemented a colocalization assay that was performed prior to each experiment. In this assay, we selected the widely expressed “housekeeping” gene *ama-1* (encoding RNA polymerase II) as a control and designed two independent sets of primary probes targeting distinct regions of its mRNA. The first probe set hybridized to the 5′ region of the transcript, while the second set targeted the 3′ region, and each was detected by distinct readout probes in separate fluorescent channels. The degree of colocalization between fluorescent spots from the two channels was used as a quantitative measure of probe specificity. Colocalization efficiency was quantified across 10 randomly selected regions pooled from 3 independent experiments, yielding a mean of 77.43% ± 4.66% (SD).

### Image processing

After image acquisition and postprocessing (**Fig. 2a, b**), fluorescent transcript spots were detected and decomposed into single-molecule signals using the Big-FISH software library^31^ (**Fig. 2a, c)**. Because our images included multiple cell types with variable nuclear sizes and morphologies, we trained a custom model tailored to our dataset using the StarDist Python library^32^. This model successfully identified DAPI-stained nuclei across diverse cell types and assigned accurate three-dimensional nuclear masks across multiple z-stacks (**Fig. 2d**).

**Figure 2.**
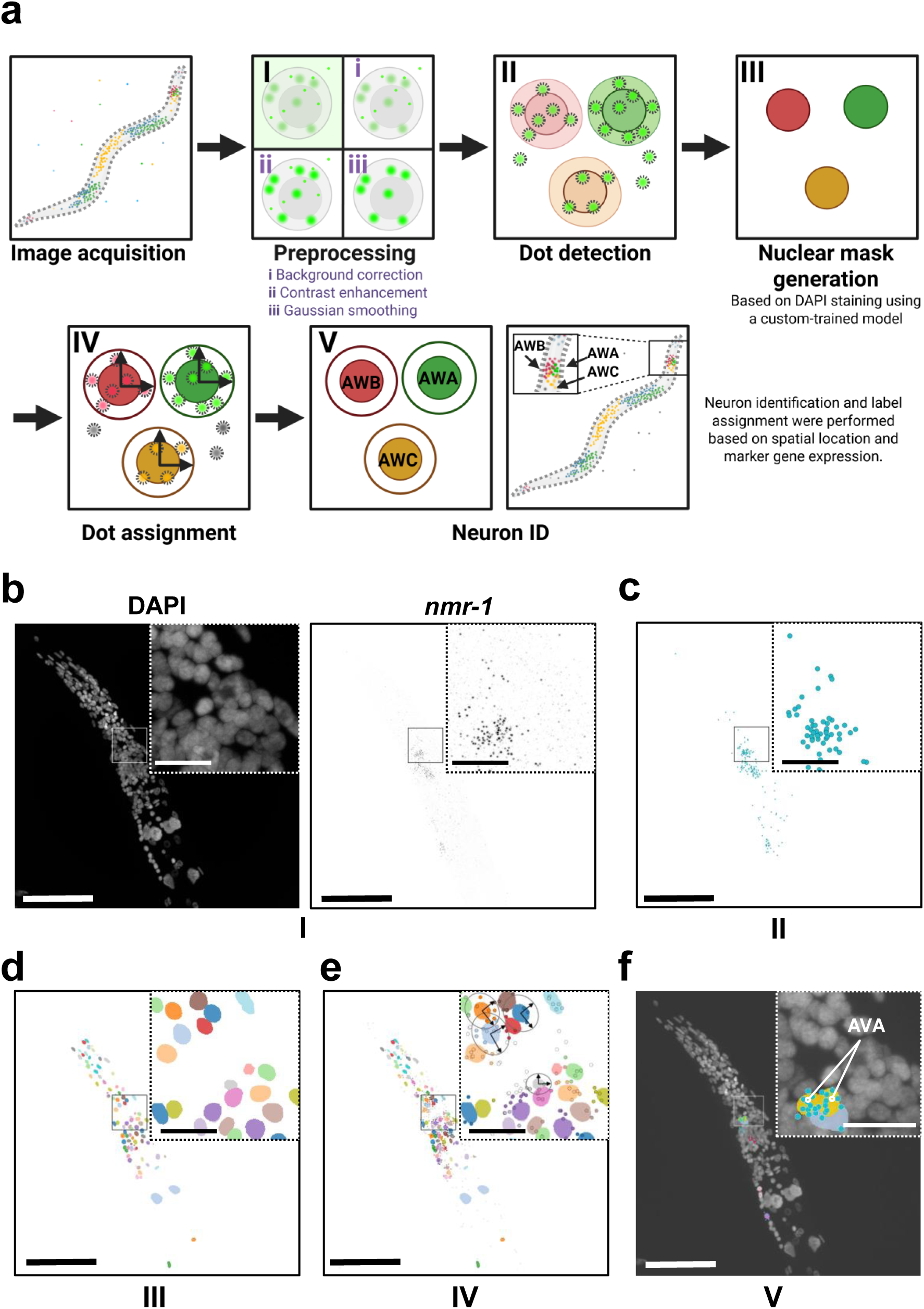
Overview of imaging and analysis. (a) Image processing workflow. (I) Acquired images were background-corrected, contrast-enhanced, and filtered with a low-pass Gaussian filter. (II) Using the Big-FISH library, smFISH signals are detected and decomposed into individual dots. (III) Nuclear segmentation is performed using the StarDist library with a custom-trained model, and each nucleus is assigned a nuclear mask. (IV) Transcript-to-cell assignment is using nuclear masks. (V) Manual neuron identification is achieved by integrating marker gene expression patterns with anatomical position, after which nuclear masks are assigned to their corresponding neuron identities. (b) Preprocessed image after background correction, contrast enhancement, and low-pass Gaussian filtering to improve signal visualization and reduce noise. DAPI was imaged at 405 nm and *nmr-1* at 640 nm. (c) Dot detection using the Big-FISH library with a manually defined threshold. (d) Nuclear segmentation performed using the StarDist library with a custom-trained model. Representative Z = 17. (e) Transcript-to-cell assignment based on nuclear masks, as described in (a-IV). Representative Z = 17. (f) Neuron identification achieved by integrating marker gene expression patterns with anatomical position, with nuclear masks assigned to the identified AVA neuron based on *nmr-1* expression, overlaid on DAPI maximum intensity projections. Scale bar, 50 μm; inset (magnified region), 10 μm.

A major challenge in our analysis was assigning detected transcripts to the correct cells. Unlike most spatial transcriptomics approaches, which rely on membrane staining for cell segmentation^15,33^, we used the nuclear masks generated from our segmentation model as the basis for transcript assignment. Detected transcripts were mapped to individual nuclei based on spatial proximity: transcripts located within a nuclear boundary were directly assigned to that nucleus, while those within the perinuclear region (defined as within twice the nuclear size, up to a maximum of 15 pixels) were probabilistically assigned using a Gaussian Mixture Model that accounted for local nuclear geometry. This strategy minimized biases related to nuclear size and shape variability and enabled more accurate transcript-to-cell mapping (**Fig. 2a, e**).

Once transcripts were assigned to specific nuclear masks, we next identified individual neuron classes to map transcriptional expression patterns and quantify transcript abundance at single-cell resolution. To achieve this, we used a panel of well-characterized marker genes with established cell-type–specific expression patterns^2,34^. By integrating marker gene expression with the anatomical position of each nucleus, we were able to distinguish distinct neuronal classes and assign corresponding neuron identities to individual nuclear masks (**Fig. 2a, f**). In males, a mean of 13.56 ± 5.68 (SD) genes were detected per neuron class among the 81 genes assayed, using a mean value of at least 1 count per class to call a detection, with each class estimate averaged across 3-9 worms. Detection was consistent with the observed cells of each neuron class: genes deemed detected for a class were observed on average in 79.6% ± 22.6% (SD) of cells, using a soft-assignment cutoff of 0.5 counts. For hermaphrodites, a mean of 8.40 ± 4.98 (SD) genes were detected per class among the 57 genes assayed, using a mean value of at least 1 count per class to call a detection, with each class estimate averaged across 2–6 worms, and detected genes were observed on average in 86.6% ± 22.2% (SD) of cells. This approach enabled accurate quantification of gene expression in intact *C. elegans* while preserving spatial context and allowing the assignment of detected transcripts to specific neuron types.

### *C. elegans* sequential smFISH approximates single-cell RNA-seq measurements

Sequential smFISH was used to generate gene expression profiles across *C. elegans* neuron classes in hermaphrodites and males. To assess transcript detection at single-cell resolution and evaluate overall data reliability, we compared sequential smFISH measurements with both single-cell (CeNGEN)^2^ and bulk RNA-seq datasets^35^. In hermaphrodites, heatmaps of averaged neuron class expression (1–4 single neurons per class) revealed distinct transcriptional signatures (**Fig. 3a**). Corresponding single-cell RNA-seq profiles from the CeNGEN hermaphrodite dataset showed similar class-specific expression patterns (**Fig. 3b**) except for some genes in the tail region, where reduced detection was observed. Equivalent analysis in males showed comparable resolution and reproducibility across neuron classes (1–4 single neurons per class; **Fig. 3c**), with expression patterns consistent with the CeNGEN male dataset^2^ (**Fig. 3d**), although overall transcript levels were slightly lower compared to CeNGEN. To assess global correspondence with sequencing data, total gene expression measured by sequential smFISH was compared with bulk RNA-seq^35^. Expression levels were strongly correlated in both hermaphrodites (2–6 worms; **Fig. 3e**) and males (3–9 worms; **Fig. 3f**). The overlap in gene–neuron detection between sequential smFISH and RNA-seq was further evaluated using Venn diagrams summarizing the shared and unique gene–neuron pairs (**Fig. 3g, j**). In hermaphrodites, 130 out of 359 gene–neuron pairs were detected by both methods, with 40 pairs identified exclusively by sequential smFISH. In males, 285 out of 884 pairs were shared between sequential smFISH and the CeNGEN dataset, with 121 pairs detected exclusively by sequential smFISH. Differences in detection sensitivity between methods may account for some of the discrepancies, as scRNA-seq datasets such as CeNGEN aggregate transcripts across many individual cells, potentially enabling detection of low-abundance gene–neuron pairs that fall below the sensitivity threshold of sequential smFISH. Quantitative comparison of single-cell expression levels across both datasets showed high concordance for neuron-class–specific genes in hermaphrodites (**Fig. 3h**) and males (**Fig. 3i**), with data points distributing along the diagonal, indicating that genes with high expression in CeNGEN correspondingly show high expression in sequential smFISH. This indicates that sequential smFISH captures single-cell RNA-seq expression patterns while preserving native spatial context at single-cell resolution.

**Figure 3.**
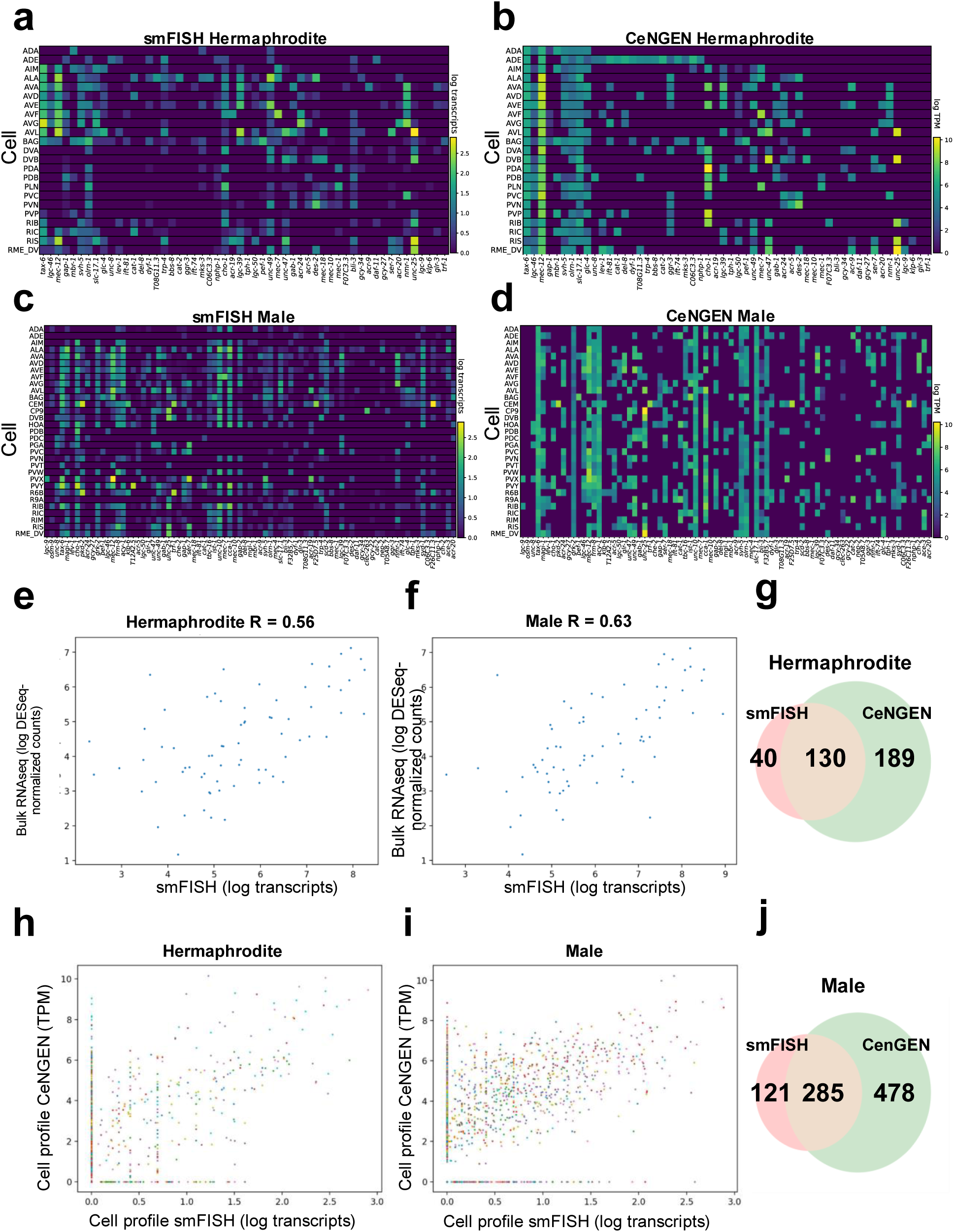
Gene expression profiles of the male and hermaphrodite neurons and comparisons to RNA-seq. (a) Heatmap of neuron class gene expression profiles detected with sequential smFISH in *C. elegans* hermaphrodite. (b) Heatmap of the corresponding single cell RNA-seq from the CeNGEN hermaphrodite dataset for comparison with (a). (c) Heatmap of neuron class gene expression profiles detected with sequential smFISH in *C. elegans* male. (d) Heatmap of the corresponding single cell RNA-seq from the CeNGEN male dataset for comparison with (c). (e) Comparison of total gene expression quantification in sequential smFISH with hermaphrodite bulk RNA-seq data. Each point represents the average total expression of a different gene in a different neuron class. (f) Comparison of total gene expression quantification in sequential smFISH with male bulk RNA-seq data. (g) Venn diagram displaying the number of gene–neuron pairs detected in sequential smFISH versus single cell RNA-seq in the CeNGEN dataset to represent the overlap in the heatmaps in (a) and (b). (h) Comparison as in (e) with CeNGEN hermaphrodite dataset. Each point represents the average expression of a specific gene in a specific neuron class, and each gene is assigned to a different color. (i) Comparison of single cell gene expression quantification in sequential smFISH with single cell RNA profiles from the CeNGEN male dataset. (j) Venn diagram as in (g) to represent the overlap in the heatmaps in (c) and (d).

### Identification of neuronal types by marker gene expression

To facilitate neuron identification in future *C. elegans* sequential smFISH experiments, we aimed to generate a curated list of marker genes that enable reliable recognition of distinct neuronal classes. We focused on neurons in the head and tail regions by curating a set of genes using publicly available databases^2,36^. Due to limitations on the gene panel size, the selected marker genes were distributed across two experiments (**Fig. 4**). Gene expression profiles from different worms imaged in each of the experiments were subsequently combined. The pipeline was optimized to ensure that cell type assignments remain consistent across rounds, making the integration of multiple smaller gene panels both feasible and reliable. Using these markers, we were able to identify up to 86 neuronal classes, including 27 that were male-specific. Neuron classes were assigned to nuclear masks whose transcript profiles matched the expected marker gene single-cell expression patterns^2^ at the correct anatomical positions^37–40^.

**Figure 4.**
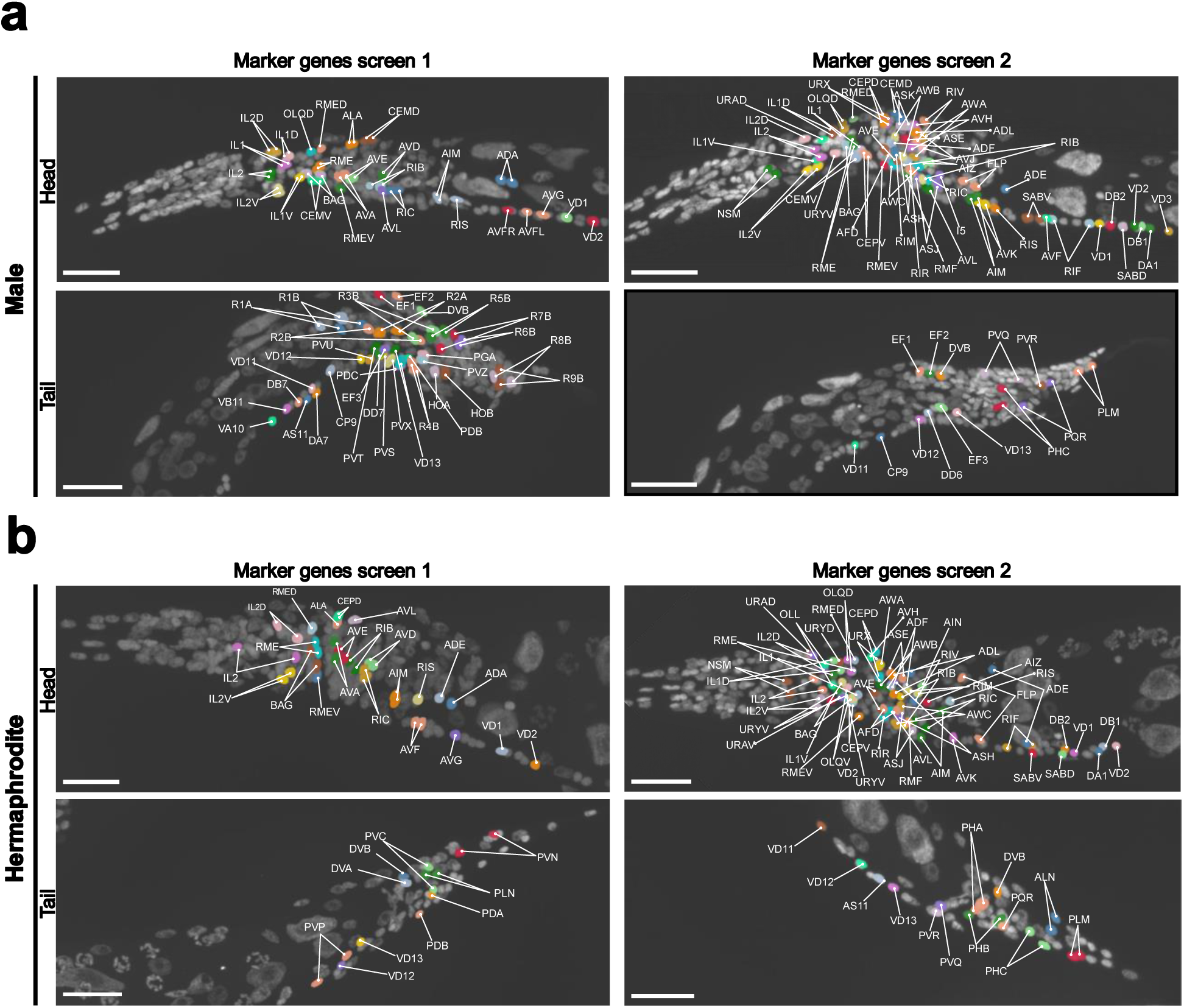
Nuclear masks of identified neurons based on marker gene expression. Neuron identification was performed across two experiments. Nuclear masks corresponding to different neuronal types are color-coded. (a) Identified neurons in the head and tail regions of the *C. elegans* adult male. (b) Identified neurons in the head and tail regions of the *C. elegans* adult hermaphrodite. Scale bar, 20 μm.

To further assist in experiment design, we defined three optimized marker gene sets that allow flexible neuron annotation depending on the desired resolution (**Fig. 5**). For each marker set, we provide the probe designs validated in this study that perform robustly (**Supplementary Table 2**). The first set, comprising 16 genes, is designed to label the maximum number of neuron classes possible, enabling the identification of up to 86 neuron classes in both hermaphrodites and males. The second and third sets are designed for region-specific annotation using the minimum number of genes necessary: a set targeting head neurons distinguishes approximately 51 neuron classes, while a set of 11 genes targeting tail neurons allows identification of up to 40 neuron classes, with 5 genes shared between both regions.

**Figure 5.**
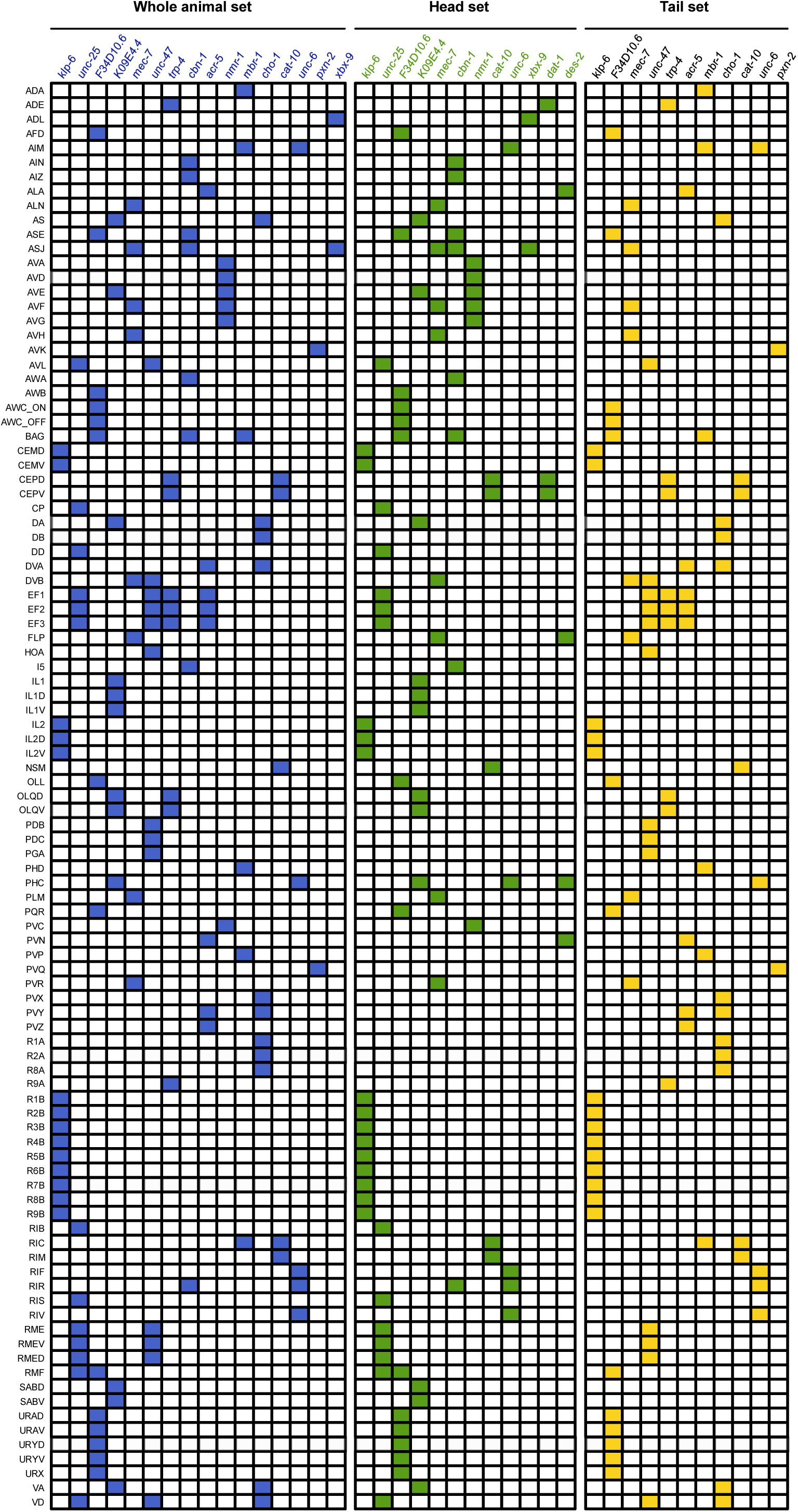
Marker gene expression table based on sequential smFISH detection for the identification of distinct neuron sets in both male and hermaphrodite adult *C. elegans*, in combination with anatomical position. Three marker gene sets were designed: one targeting neurons in both head and tail regions (blue), one specific to head-region neurons (green), and one specific to tail-region neurons (yellow).

### Sequential smFISH reveals sexually dimorphic expression of genes in *C. elegans*

To demonstrate the ability of our method to accurately detect gene expression patterns at the single-cell level, we used publicly available datasets^2,35,41–43^ (**Supplementary Table 1**) to compile a list of 100 candidate genes predicted to exhibit sexually dimorphic expression (**Supplementary Fig. 2**). Of these, 75 genes were successfully imaged in our assay. During our screen, we identified several genes with clear sex-specific expression patterns. For instance, *paml-2* (F25D7.5), F26C11.3, and *trf-1* were expressed in male-specific sensory ray B neurons and CEM cephalic neurons, with *paml-2* (F25D7.5) also showing shared expression in the sex-common PVP neuron (**Fig. 6a, b**). *trf-1* encodes a TRAF-like adaptor protein and, together with *trf-2*, is required in male extracellular vesicle (EV)–releasing neurons for mating regulated by the polycystins (LOV-1 and PKD-2)^44^, ciliary sensory proteins that detect mating cues. *paml-2* (F25D7.5) and F26C11.3 are similarly expressed in male EV-releasing neurons involved in sensory detection during mating^44^. We also observed *cat-2* expression (encoding tyrosine hydroxylase, the rate-limiting enzyme in dopamine biosynthesis)^45,46^ in male ray A sensory neurons, as well as shared expression in CEP and ADE mechanosensory neurons. These expression patterns are consistent with previous reports based on cell purification and reporter gene analysis^2,43–45^ and our results directly confirm their sexually dimorphic expression at single-cell resolution in intact animals.

**Figure 6.**
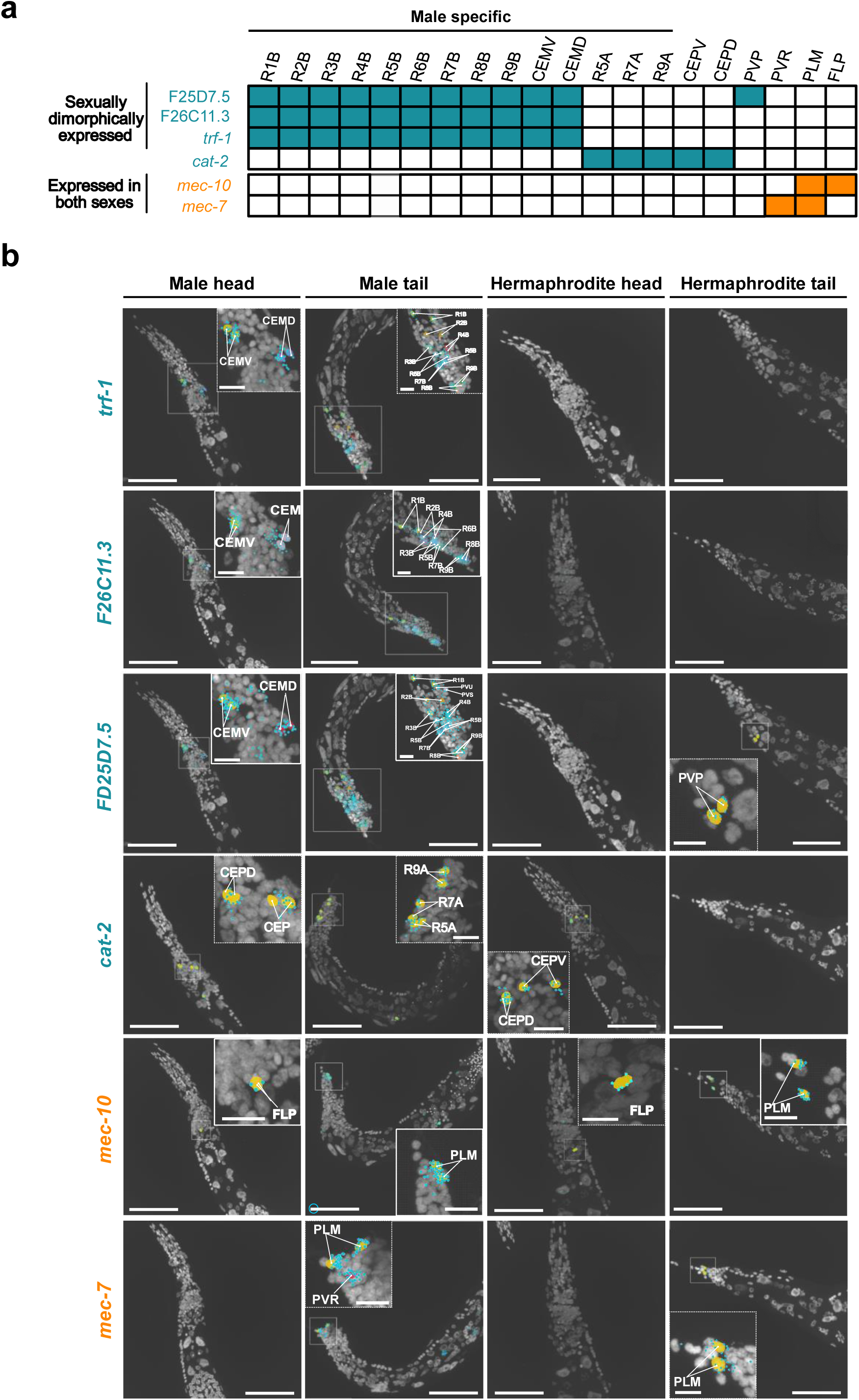
Sexually dimorphic gene expression. (a) Summary table of gene expression patterns showing sexually dimorphic genes (blue) and genes with shared expression across a limited number of neuron types (orange). (b) Expression patterns of the genes listed in (a) in the head and tail regions of *C. elegans* adult males and hermaphrodites. Nuclear masks corresponding to identified neuron types are overlaid on DAPI maximum intensity projections. Scale bar, 50 μm; inset, 10 μm.

In addition to sexually dimorphic genes, we detected transcripts with highly restricted expression patterns, suggesting potential utility for the generation of genetic drivers specific to individual neuron classes. For example, *mec-7* showed strong expression in the PLM mechanosensory neurons in the tail (**Fig. 6b)** and in the PVR interneuron. Similarly, *mec-10* was highly expressed in the PLM neurons, as well as in the FLP neurons of the head region.

Together, these results demonstrate that our method enables accurate, high-resolution mapping of complex gene expression patterns across both sexes of *C. elegans*, providing a robust framework for identifying neuron-type-specific genes and for guiding future genetic and functional studies of neural diversity.

## Discussion

### Multiplex spatial transcriptomics in *C. elegans*

Here, we present a spatial transcriptomics approach for anatomically intact *C. elegans* that combines multiplexed detection of mRNA transcripts, computational assignment of mRNA transcripts to cell nuclei, and marker gene panels for neuronal cell-type identification. The method allows for the spatially resolved gene expression analysis in multiple individual worms per experiment without the need for sectioning. Using targeted gene panels, we imaged a total of 40 genes per experiment and benchmarked the results against single-cell and bulk RNA-seq datasets^2,35^, showing that our approach can complement sequencing-based resources with data from the preserved spatial context of whole-mount worm preparations.

### Probabilistic assignment and curated markers enable robust neuron identification despite segmentation challenges

An important feature of the method is the assignment of mRNA transcripts to cells based on DAPI-derived nuclear masks, which can be readily recorded for whole-mount *C. elegans* preparations. This robust nuclear-dye segmentation approach assigns transcripts within a fixed distance of the nucleus, and thereby mainly captures cell body transcript signals. Similar to previously described methods^47^, it leaves mRNAs located to neuronal projections or other elongated, non-spherical cellular regions unassigned. Developing methods to robustly stain membranes in worms or otherwise improve segmentation could help recover these signals and provide more complete cell-level transcriptomic data.

In the data shown here, overall gene expression profiles closely approximate both bulk and single-cell RNA-seq measurements^2,35^. Differences between our data and reference scRNA-seq datasets can reflect both method differences and downstream processing choices, including segmentation, filtering, and transcript-to-cell identity mapping^48–50^. When we compared our data to published single-cell profiles from the hermaphrodite tail neurons^2^, a subset of expected genes showed reduced or no detection (**Fig. 3a, b**). This discrepancy likely reflects low mRNA abundance in this region, together with the limitations of nuclear-mask-based assignment. When transcript density is sparse, probabilistic assignment becomes less reliable, and transcripts localized to neuronal projections fall outside the assignment zone, resulting in under-detection of transcripts.

### A scalable framework for reproducible neuronal mapping

A major goal of this method was to make neuron identification reproducible and accessible. Manual assignment of neurons is labor-intensive, and screening for neuron-specific marker genes requires substantial effort and resources. To streamline neuron identification, we provide a curated marker gene panel that enables reliable identification of most neuron classes within our workflow. To build this panel, we first evaluated a set of 13 marker genes that had previously been shown to successfully identify neuronal classes^36^. However, only 9 of the 13 markers proved informative in our hands, as 4 were too broadly expressed to reliably distinguish specific neuron types. With this initial set alone, the number of confidently identifiable neuron classes remained limited. We therefore performed an additional screen of 114 genes curated from the CeNGEN datasets^2^, which allowed us to identify markers for additional neuronal classes. Combined with the 9 validated markers from the initial set, our final gene panel enables identification of a total of 86 neuron classes, including 27 male-specific classes (**Figs. 4 and 5**). Because marker genes must be included in every imaging round, we then optimized a minimal marker panel that balances discriminative power with the 40-gene capacity of our current implementation, while preserving sufficient panel capacity for biological targets of interest. Experiments with higher multiplexing could adopt more scalable strategies, such as designing combinatorial marker sets whose unique expression patterns label nearly all neurons without limiting the number of genes available for biological measurements. In addition, users can select a subset of markers from the list provided here to prioritize specific neuron types and to further maximize capacity for genes of interest.

### Sequential smFISH detects sexually dimorphic gene expression patterns

During our screen, we confirmed expression of several genes in ray and CEM neurons, previously characterized by cell purification or reporter gene analysis^2,43–45^. For these and other genes, we now provide additional data about their sexually dimorphic expression. As expected, *cat-2* (encoding tyrosine hydroxylase, the rate-limiting enzyme in dopamine biosynthesis)^45,46^ was detected in male sensory ray neurons and head CEM neurons. We also observed male-specific expression of *trf-1* in ray neurons, confirming previous results from transcriptomic studies. *trf-1* encodes a TRAF-like adaptor protein and, together with *trf-2*, is required in male extracellular vesicle (EV)–releasing neurons for mating regulated by the polycystins (LOV-1 and PKD-2)^44^, ciliary sensory proteins that detect mating cues. In addition, we observed male-specific expression of *paml-2* (F25D7.5) and F26C11.3 in ray and CEM neurons, both of which were identified in extracellular vesicle enriched neurons. Both genes are expressed in male EV-releasing neurons involved in sensory detection during mating^44^. These are examples of how the use of sequential smFISH allowed us to map multiple genes simultaneously at single-cell resolution, revealing spatial patterns that would be missed by bulk transcriptomic approaches.

### Whole-organism spatial transcriptomics for cross-tissue signaling and cellular asymmetry

Many biological processes in *C. elegans* involve long-range communication between neurons and peripheral organs, including neuroendocrine and neuropeptidergic signaling pathways that coordinate metabolism, development, and stress responses, yet such interactions must currently be inferred indirectly from genetics or reporter-based experiments. For example, neuronal insulin/IGF signaling influences intestinal metabolism and lifespan^51^, serotonergic neurons trigger secretion of the neuropeptide FLP-7 to regulate fat metabolism^52^, gut-brain axis peptide signals modulate neuronal control of metabolic programs^53^, and the decision between dauer formation and reproductive development depends on insulin-like peptides from neurons signaling to the intestine^54^. Therefore, whole-organism spatial transcriptomics provides a unique opportunity to directly map such gene expression interactions at single-cell resolution. Besides inter-tissue signaling, this approach also allows for the direct comparison of transcriptional states across individuals and between asymmetric neuronal pairs that are morphologically identical yet functionally distinct, such as the bilaterally asymmetric ASE neurons (ASEL/ASER) and olfactory AWC neurons (AWC_ON_/AWC_OFF_)^55–58^.

In summary, our method provides a scalable and reproducible framework for whole-animal spatial transcriptomics in *C. elegans*. By combining smFISH with optimized sample preparation and curated marker sets, it enables high-resolution, spatially resolved profiling of multiple neuronal classes in a single experiment. While challenges such as segmentation limitations and probabilistic transcript assignment remain, these can be addressed in future work through improved labeling and analysis strategies. Overall, this approach opens the door to comprehensive studies of gene expression, neural circuitry, and organism-wide signaling.

## Methods

### Strains and culture conditions

*C. elegans* strains N2 (Bristol) wild-type and CB4088 *him-5(e1490)* (segregating about one-third males) were used. Strains were maintained under standard laboratory conditions using Nematode Growth Medium (NGM) plates with *E. coli* OP50 as the food source as described by Brenner (1974)^59^.

### Primary probe design

Probe sets were generated as previously described^60^. An initial gene pool including probes targeting exons within the coding regions of 114 unique transcripts comprising 100 target genes, 13 marker genes, and 1 housekeeping gene as control was designed for the initial screening of differentially expressed genes. Due to experimental setup constraints, the assay is limited to two fluorescent channels. With 20 hybridization rounds, a maximum of 25 target genes and 15 marker/housekeeping control genes can be screened per experiment. For neuron identification downstream, the 13 marker genes and 1 control gene (targeted by two distinct readouts for colocalization assay) were included in every experiment, resulting in a total of 40 genes per experiment. An additional gene pool including 55 marker genes and 1 housekeeping gene as control was prepared for additional marker genes screening. Each probe was 35 nucleotides in length, with successive probes spaced by 2 nucleotides, and a GC content ranging from 40–65%. A local BLAST query was performed for each probe against the *C. elegans* transcriptome to assess specificity. Probes with BLAST hits of 17 nucleotides or longer to sequences other than the intended target were considered off-target and removed. To further minimize cross-hybridization between probe sets, a local BLAST database of all probe sequences was constructed, and any probes with 17-nucleotide or longer matches were removed from the larger probe set. Target genes that did not yield a minimum of 24 probes were excluded.

### Readout probe design

Probes were designed following previously described methods^61^. Random 15-nt oligonucleotide sequences with a GC content of 40–60% were generated, and each sequence was screened for specificity using a local BLAST search against the *C. elegans* transcriptome. Any probe containing a contiguous 10-nt match was excluded.

### Primary probe pools amplification

Primary probes were obtained as oligoarray complex pools from Twist Bioscience and prepared as previously described, with minor modifications^60^. Briefly, the probe sequences were amplified from the oligo complex pool using a limited number of PCR cycles, during which a T7 promoter site was appended to the sequences. The resulting PCR products were purified with the QIAquick PCR Purification Kit (Qiagen, 2 10) according to the manufacturer’s instructions. These purified PCR products served as templates for in vitro transcription (NEB, E2050S) and were supplemented with RNasin RNase Inhibitor (Promega, N261A) and Pyrophosphatase (NEB, M0361S).

The transcribed RNA was subsequently reverse transcribed using Maxima H Minus Reverse Transcriptase (Thermo Fisher Scientific, EP7051) along with RNasin RNase Inhibitor and Pyrophosphatase. Following reverse transcription, the RNA templates and forward primers were removed via alkaline hydrolysis using 250 mM NaOH at 65 °C for 30 minutes, after which the reaction was neutralized with acetic acid at a final concentration of 250 mM. The resulting DNA probes were precipitated using a 30:1 solution of ethanol to sodium acetate at −20 ° for 30 minutes and resuspended in water.

Finally, the probes were purified using SPRIselect beads (Beckman Coulter, B23318) with minor modifications to the manufacturer’s instructions. In brief, DNA strands were captured with a 2× SPRI volume relative to the starting material, washed with 80% EtOH on a magnetic rack, and eluted in water. Purified probes were stored at −20 ° until use.

### Readout probe conjugation

Readout probes, consisting of 5′ amine-modified single-stranded DNA (15-nt), were obtained from Integrated DNA Technologies (IDT). Dye conjugation of the readout probes was carried out as previously described, with minor modifications^62^. In brief, 5 nmol of amine-modified oligonucleotides were resuspended in 0.5 M sodium bicarbonate buffer to a final concentration of 250 μ. To this solution, 0.5 mg of Alexa luor S ester (Thermo Fisher Scientific, A20006) or Cy3B NHS ester (Cytiva, PA63101) was added, and the reaction mixture was incubated overnight at 37 °C in the dark. Following the reaction, acetic acid was added in a molar amount equivalent to the sodium bicarbonate to quench the reaction. The dye-conjugated ssDNA probes were then precipitated using a 30:1 solution of ethanol to sodium acetate at −20 ° for 30 minutes, resuspended in water and purified via HPLC. Probes were lyophilized and resuspended in water, quantified using a Nanodrop, and diluted to a 1 mM working stock. All readout probes were stored at −20 ° until further use.

### *C*. *elegans* in situ hybridization

All procedures were performed using RNase-free consumables. Unless otherwise specified, solution exchanges were accompanied by centrifugation at 300 g for 1 minute in a benchtop microcentrifuge.

### Collection and fixation

Live *C. elegans* were collected by gently adding RNase-free H₂O (Thermo Fisher Scientific, 10977-015) to the culture plate, avoiding disruption of the bacterial lawn. Animals were washed three times with 1 mL RNase-free H₂O, repeating until the supernatant was clear. After the final wash, worms were chilled at −20 ° for 1 minute, and residual water was removed.

Fixation was performed for exactly 45 minutes at room temperature (RT) in 1 mL freshly prepared 4% paraformaldehyde (PFA; 16% stock, Thermo Fisher Scientific, 28908) diluted in 1× PBS (Ambion, AM9625). Fixed worms were washed once rapidly in 1× PBST (1× PBS containing 0.01% Triton X-100; Millipore Sigma, 93443), followed by two 15-minute washes in fresh 1× PBST at RT. The pellet was resuspended in 100% methanol (VWR, BDH1135-4LG) and stored overnight at −20 ° with tubes placed horizontally.

### Rehydration, redox treatment and clearing

All incubation steps were performed while rocking. Without removing methanol, 1× PBST was added to the tube to reach a final 75% MeOH and mixed for 30 minutes at RT. Subsequently, 1× PBST (final 50% MeOH) was added and mixed for another 30 minutes. After centrifugation, the supernatant was removed and replaced with pre-mixed 25% MeOH + 1× PBST, followed by a 30-minute incubation. Samples were centrifuged again, and the solution was replaced with 1× PBST for 30 minutes.

Worms were rinsed once in BOT buffer (1× borate buffer; Electron Microscopy Sciences, 11455-90; 0.01% Triton X-100) and treated for 30 minutes at RT with freshly prepared 100 mM TCEP (Thermo Fisher Scientific, 77720) in BOT. After a quick rinse in BOT, samples were incubated for 10 minutes at RT in 0.3% H₂O₂ (Honeywell, 216763) prepared in BOT. Samples were subsequently washed once in BOT and once in PBST.

Clearing was performed for 45 minutes at 37 °C in 8% SDS solution consisting of 10% SDS (Thermo Fisher Scientific, AM9822), 1× PBS (Ambion, AM9625), RNase-free H₂O, and 0.1% Triton X-100 (Millipore Sigma, 93443). Tubes were placed horizontally during incubation. After clearing, samples were washed once briefly with PBST and twice for 15 minutes in fresh 1× PBST.

### Cuticle digestion and postfixation

For cuticle digestion, Collagenase Type IV (Millipore Sigma, C5138) stock solution (1 kU/mL) previously prepared in buffer containing 50 mM Tris-HCl pH 8.0 (Thermo Fisher Scientific, 15568-025), 150 mM NaCl (Thermo Fisher Scientific, AM9760G), and 1 mM CaCl₂ (Millipore Sigma, 21115) was diluted in collagenase dilution buffer (150 mM Tris-HCl pH 8.0, 850 mM NaCl, 80 mM CaCl₂) for a final concentration of 0.2 kU/mL. Samples were incubated at RT for exactly 10 minutes, followed by four rapid washes with 1× PBST.

Samples were post-fixed for 30 minutes at RT in 7.5 mM BS(PEG)5 (PEGylated bis(sulfosuccinimidyl)suberate) (Thermo Fisher Scientific, A35396) freshly prepared in 1× PBS. After fixation, the solution was replaced four times at 30-minute intervals with 100 mM N-(propionyloxy)succinimide (Millipore Sigma, 93535-1G) in 1× PBS, prepared immediately before use. Samples were then rinsed twice with wash buffer (40% formamide, 2× SSC, 0.1% Triton X-100).

### Primary probe hybridization

A 1.67× hybridization buffer (30% formamide; Ambion, AM9344; 4× SSC, Thermo Fisher Scientific, 15557-044; 16.7% w/v dextran sulfate; Millipore Sigma, D8906) was diluted to 1× for use. Animals were pre-incubated in 100 µL of hybridization buffer without probes for 30 minutes at 37 °C. Worms were then centrifuged at 6,000 g for 2 minutes, and the solution was replaced with 100 µL of hybridization buffer containing 10 nM primary smFISH probes (IDT or Twist pool, NA) and 1 µM LNA-modified poly-T₁₈ oligonucleotide with a 5′ acrydite modification (IDT, NA). Samples were incubated with the hybridization solution for 48–72 hours at 37 °C.

Following hybridization, animals were washed four times quickly with 4× SSCT (4× SSC + 0.1% Triton X-100), followed by a 30–60 minute wash in 30% formamide wash buffer (30% formamide, 2× SSC, 0.1% Triton X-100) at RT. After the formamide wash, samples were washed three times with 4× SSC for 15 minutes each at RT.

### Hydrogel embedding and proteolytic digestion

Samples were incubated overnight at 4 °C with chilled monomer solution (4% acrylamide/0.2%, ′-methylenebisacrylamide premix; Bio-Rad, 1610154; in PBS).

For gelation, a fresh monomer solution was prepared on ice with the addition of VA-044 initiator (25% w/v stock; TCI, A3012) to a final 2% w/v. The solution was bubbled with nitrogen on ice for 15 minutes to remove oxygen. Worms were then mounted onto functionalized coverslips and covered with the activated monomer solution. Samples were overlaid with a Gel-slick-coated coverslip (Lonza, 50640) and polymerized for 4 hours at 37 °C in a humidified chamber.

After polymerization, the top coverslip was gently removed and a microfluidic chamber was placed on top of the gels. Gels were digested for 48 hours at 37 °C in Proteinase K buffer (50 mM Tris-HCl pH 8.0, 1 mM EDTA, 500 mM NaCl, 1% SDS, 0.5% Triton X-100) containing Proteinase K (NEB, P8107S), diluted 1:100 from an 800 unit/mL stock, yielding a final concentration of 8 unit/mL, followed by 2–3 washes in 1× PBST.

### Readout probe hybridization

Readout probes (50–100 nM; IDT or self-conjugated, NA) were hybridized for 30 minutes at RT in 10% ethylene carbonate (Millipore Sigma, E26258) hybridization buffer (2× SSC, 0.1% Triton ×0, 16.7% low-molecular-weight dextran sulfate; Millipore Sigma, D4911). Hybridized gels were washed three times in 4× SSCT, followed by a 15-minute wash in 10% formamide wash buffer (10% formamide, 2× SSC, 0.1% Triton X-100), and then washed three times with 4× SSCT.

### Counterstaining and anti-bleaching buffer

Nuclei were stained for 5 minutes with 3 μg/m DAPI (Millipore Sigma, D8417) in 4× SSC, followed by three washes in 4× SSCT. Samples were equilibrated immediately before imaging in anti-bleaching buffer containing 50 mM Tris-HCl pH 8.0, 300 mM NaCl, 0.8% D-glucose (Millipore Sigma, G7528), 0.5 mg/mL glucose oxidase (Millipore Sigma, G2133), 30 units/mg catalase (Millipore Sigma, C3155-50MG), and 3 mM Trolox (Millipore Sigma, 238813).

### Sequential Imaging

The imaging routine was performed as previously described with minor modifications^61^. Briefly, the flow cell containing the sample was connected to an automated fluidics system operating at a flow rate of 250 μ /min. The region of interest (I) was identified based on nuclear signals stained with 3 μg/m DAPI in 2× SS.

For serial smFISH experiments, each hybridization buffer contained two unique readout sequences conjugated to Alexa Fluor 647 or Cy3b (100 nM) in 10% ethylene carbonate (Millipore Sigma, E26258) hybridization buffer (2× SSC, 0.1% Triton X-100, 16.7% low-molecular-weight dextran sulfate; Millipore Sigma, D4911). A total of 100 μ of hybridization buffer including a repeat round for control round 1, was pipetted into a 96-well plate. During each serial hybridization, the automated sampler aspirated 100 μ of hybridization buffer from the designated well through a multichannel fluidic valve (IDEX Health & Science, EZ1213-820-4) to the flow cell (∼30 μ re uired) using a syringe pump (amilton ompany, 63133-01), followed by a 25-minute incubation at room temperature.

After hybridization, the sample was washed with 00 μ of 10% wash buffer (10% formamide and 0.1% Triton X-100 in 2× SSC) over 6 minutes to remove excess readout probes and non-specific binding, then rinsed with approximately 200 μ of × SS. uclear staining was refreshed with DAPI solution (3 μg/m in 2× SS) for ∼30 seconds. An anti-bleaching buffer was then flowed through the sample.

Imaging was conducted on a Leica DMi8 microscope equipped with a confocal scanner unit (Andor, Dragonfly 200), an sCMOS camera (Andor, Zyla 4.2 Plus), a 63× oil objective lens (Leica, 1.40 NA), and a motorized stage (ASI, MS2000). High-power laser engine and filter sets from Andor were used to ac uire snapshots with 1 μm z-steps over nine z-slices per field.

Following imaging, a stripping buffer (55% formamide and 0.1% Triton X-100 in 2× SSC) was applied to the sample for 1 minute, followed by a 3-minute incubation, and this procedure was repeated twice more. The sample was then rinsed with 4× SSC. These steps of serial hybridization, imaging, and probe stripping were repeated until the desired number of rounds was completed. After the final stripping step, an additional round of imaging was performed to verify the efficiency of probe removal. Integration of the automated fluidics system with image acquisition was controlled using a custom-written script in μ anager.

### Image processing

Image stacks were subjected to background correction using a Gaussian-based subtraction followed by gamma adjustment. For each fluorescence channel, a Gaussian blur (σ = 3; E size = 3) was applied to estimate the local background intensity, which was then subtracted from the original image. Negative pixel values were clipped to zero. The corrected image was subsequently contrast-enhanced using a gamma correction (γ = 1.) to improve visualization of fluorescent signals. After background correction, a low-pass aussian filter (σ = 1, kernel size = 3) was applied to reduce noise while preserving spatial features relevant for single-molecule detection.

### DAPI-based alignment

To correct drift across hybridization cycles, consecutive image stacks were aligned to the reference hybridization (initial hybridization round) using the DAPI channel. For each image stack, the DAPI channel was extracted and maximum-intensity projection was generated along the z-axis. Translational offsets relative to the reference DAPI projection were calculated using phase correlation. The resulting x/y shift was applied across all z-slices and fluorescence channels of the moving stack to generate a corrected, aligned image.

### Dot detection

Dot detection and spot decomposition were performed using the Big-FISH library^31^. Detection was carried out on preprocessed images corrected for background and smoothed with a low-pass Gaussian filter. A manual intensity threshold was selected for each experiment based on visual inspection to identify spot fluorescence signals. Voxel size was estimated as (z, y, x) = (1000, 103, 103) nm and the expected spot radius as (1050, 300, 300) nm.

### Colocalization Analysis

Two sets of probes targeting the control gene *ama-1* were hybridized. The first set targeted the 5′ half of the mRNA transcript and was detected using Alexa Fluor 647–conjugated readout probes, while the second set targeted the 3′ half of the transcript and was detected using y3b-conjugated readout probes. Images from each channel were preprocessed and dots were detected independently. For each dot in the first dataset, the Euclidean distance to all dots in the second dataset was computed using a nearest-neighbor search. A dot in the first set was considered colocalized if its nearest neighbor in the second set fell within a 2-pixel radius. Colocalization efficiency was then calculated as the fraction of dots in the first channel that had a corresponding dot in the second channel. This metric was used to assess the specificity and consistency of primary probe labeling.

### Nuclear segmentation

Nuclear segmentation was performed using the StarDist library^32^. A custom 3D nucleus detection model was trained using 122 manually annotated training images (400 × 400 × 12 voxels) and applied to the DAPI channel from our image stacks for 3D nuclear segmentation.

### Neuron annotation

Neuron annotation was performed manually using *C. elegans* nervous system anatomical positions and the gene expression patterns of marker genes (Table 1). For each sample, all hybridization cycles were compiled into a single HDF5 file and visualized using a custom graphical user interface (GUI) for annotation^63^. Using anatomical drawings^37–40^, individual neurons were identified for each marker gene based on a combination of single-cell gene expression patterns^2^ and anatomical position, and an XYZ coordinate label was manually placed at the center of each neuron’s nucleus. A second round of annotation was performed to corroborate annotation accuracy; neurons for which identification was uncertain were either not annotated or were excluded from further analysis. The finalized coordinates were subsequently extracted and assigned to neuron-specific masks obtained from the 3D nuclear segmentation step, enabling integration of gene expression and anatomical identity for downstream analyses.

### Dot assignment

Detected transcripts were hard-assigned to cells if within the boundary of a nuclear mask, not assigned if beyond a threshold distance from any mask, and probabilistically soft-assigned to a cell otherwise. This approach assumes that larger nuclei possess larger cell boundaries and that probabilistic soft-assignment when there are multiple nuclei nearby should be weighted according to nuclear mask size and morphology within the 2D plane. For each z-stack, each nuclear mask was fitted with an ellipse by obtaining the first and second principal components of the pixels in the mask in that z-stack using the k-spaces package^64^, as shown in **Fig. 2**. The major and minor axes lengths were determined by doubling the length and width of the mask measured along those respective axes, producing an ellipse twice the size of the mask with an orientation determined by nucleus shape. This scheme encodes two assumptions: mRNAs found within a nuclear mask are assumed to be located in that cell, and mRNAs outside nuclear masks are assumed to be normally distributed from the nuclear center with a covariance defined by nuclear morphology. Specifically, ½ of the long diameter and ½ of the short diameter of the ellipse were used as standard deviations for a 2-dimensional normal distribution. Transcripts within this boundary were considered for assignment, while transcripts outside the ellipse were assumed not to belong to any nucleus and were left unassigned. Transcripts were then soft-assigned (with assignment probabilities summing to 1) to eligible nuclei by computing the expected probability of the transcript belonging to each nucleus within a Gaussian Mixture Model (GMM) framework using the k-spaces package^64^. Mixture component weights for the nuclei in this model were √(½ long axis length × ½ short axis length) to cancel out the bias against larger nuclear masks that would otherwise arise in a GMM with equal mixture component weights. Transcripts within the elliptical boundary for only one mask were assigned to that nucleus as a result of the framework. Transcript counts for each cell were summed across z-stacks to produce cell × gene matrices.

### Cell × gene matrices for analysis

Probabilistically soft-assigned transcript counts for each cell were summed across z-stacks to produce cell × gene matrices. To produce neuron class gene expression profiles, counts were averaged across worms per class. Marker genes were imaged in all 3 pools, while non-marker genes were only imaged in one pool each. Each pool contained 2–3 worms, and the imaging of one gene pool failed for hermaphrodites and was therefore excluded. This meant that each gene in the neuron class profile was the average of 1–4 single cells (depending on the number of cells in that class) over 3–9 worms (3 for non-marker genes but 9 for marker genes imaged in each pool) for males and 2–6 worms for hermaphrodites. For total worm gene expression profiles compared to bulk RNA-seq data, the sum over the entire cell × gene matrices was averaged across worms, accounting for the fact that non-marker genes were only imaged in one pool each.

### Comparison to CeNGEN and bulk RNA-seq data

CeNGEN neuron class profiles were obtained from the CeNGEN^2^ cell profile at https://cengen.shinyapps.io/CengenApp/ and https://cengen.shinyapps.io/male/ with the threshold “unfiltered.” Preprocessed bulk A-seq profiles were obtained from Kim et al. (2016)^35^. The M4 and M5 columns of their data were averaged to produce the male bulk profile and the H4 and H5 columns were averaged to produce the hermaphrodite profile.

## Supplementary figures

**Supplementary Table 1.** Summary of public RNA-seq datasets analyzed for sequential smFISH target gene selection. RNA-seq datasets from publicly available sources were examined to identify candidate differentially expressed genes between male and hermaphrodite *C. elegans*. Kim B. et al., (2016) dataset was partitioned into 27 co-expression modules (Mod), each representing a distinct gene expression program, from which Mods were utilized. The resulting candidate gene list was used to design the sequential smFISH probe pool.

| DATABASE | Gene set used | Number of genes | Article title |
| --- | --- | --- | --- |
| Ebbing A. et al., (2018) | Male tail enriched | 123 | Spatial Transcriptomics of *C. elegans* Males and Hermaphrodites Identifies Sex-Specific Differences in Gene Expression Patterns |
| Kaletsky R. et al., (2018) | Neuron enriched | 867 | Transcriptome analysis of adult *Caenorhabditis elegans* cells reveals tissue specific gene and isoform expression |
| Spencer W.C. et al., (2011) | Neuron enriched | 547 | A spatial and temporal map of *C. elegans* gene expression |
| Kim B. et al., (2016) | Mod 1,2,3 | 7101 | Gene function prediction based on developmental transcriptomes of the two sexes in *C. elegans* |
| Kim B. et al., (2016) | Mod 1, 2 | 3628 | Gene function prediction based on developmental transcriptomes of the two sexes in *C. elegans* |
| Kim B. et al., (2016) | Male enriched | 1751 | Gene function prediction based on developmental transcriptomes of the two sexes in *C. elegans* |
| Kim B. et al., (2016) | Mod 1,2,3 GO analysis | 123 | Gene function prediction based on developmental transcriptomes of the two sexes in *C. elegans* |

**Supplementary Figure 1.**
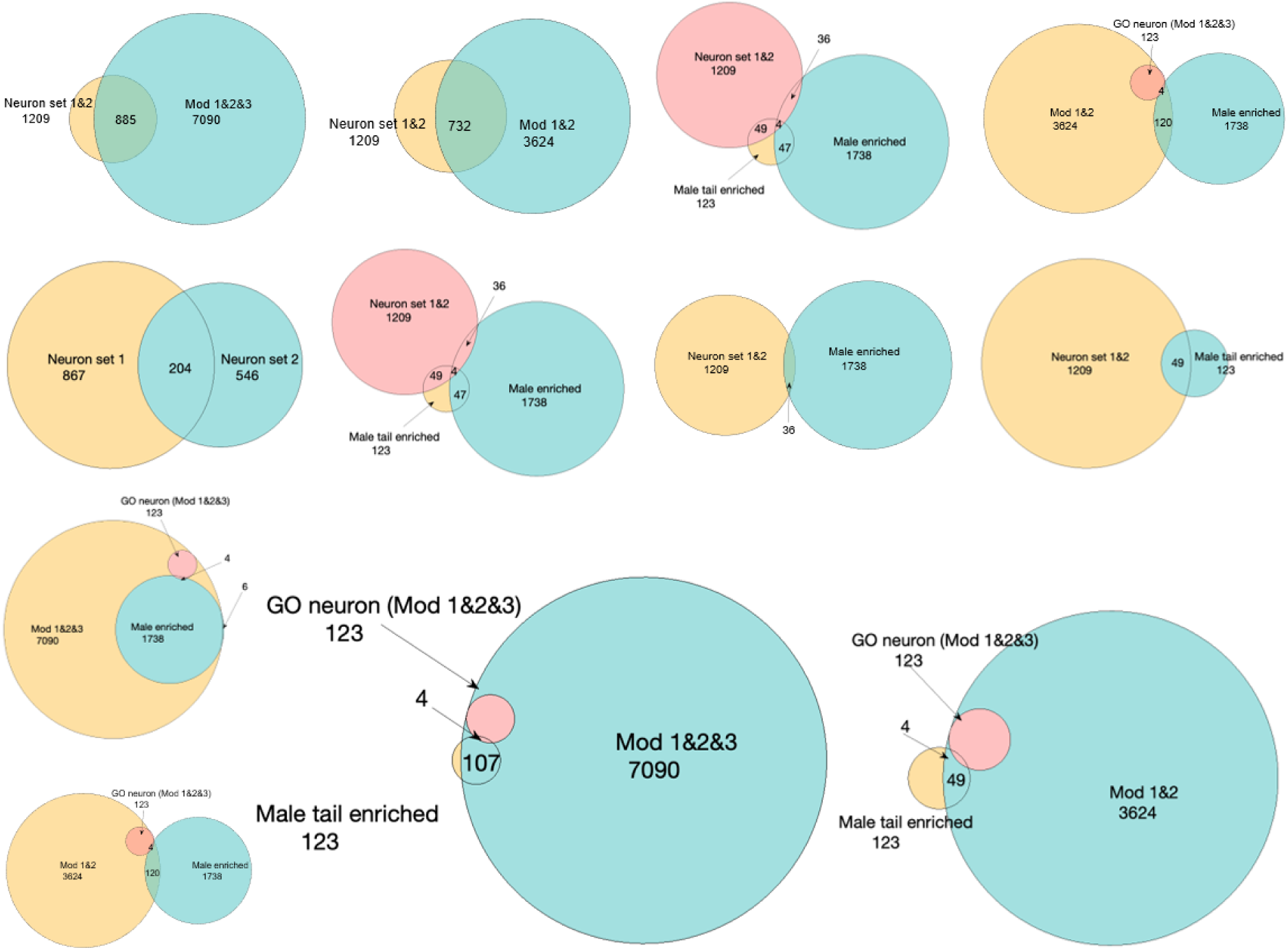
Summary of public RNA-seq datasets analyzed for sequential smFISH target gene selection. RNA-seq datasets from publicly available sources were examined to identify candidate differentially expressed genes between male and hermaphrodite *C. elegans*. Kim B. et al., (2016) dataset was partitioned into 27 co-expression modules (Mod), each representing a distinct gene expression program, from which Mods were utilized.

## Acknowledgments

We thank Stephanie Nava and Wilber Palma for their initial contributions to this work. We are grateful to Alessandro Groaz, Elsy Buitrago, Hillel Schwartz and James Tan for their discussions and suggestions. We also thank Diana Huynh and Prof. Tsui-Fen Chou for providing NIH 3T3 cells for experimental tests. This work was supported by: R01 NS113119 (P.W.S., A.D.T.S.), Tianqiao and Chrissy Chen Institute for Neuroscience senior postdoc fellowship (X.W.), Tianqiao and Chrissy Chen Institute for Neuroscience postdoc innovator grant (X.W.) and Tianqiao and Chrissy Chen Institute for Neuroscience Systems Neuroscience Awards (P.W.S.). P.W.S. is Bren Professor of Biology. The *Caenorhabditis* Genetics Center provided some strains.

## Author contributions

P.W.S. X.W., C.T. and J.D.A. conceived the experiments. J.D.A., X.W. and C.T. performed the experiments. N.M., J.D.A. and F.P. developed the data analysis pipeline. L.C. and P.W.S. supervised the project. All authors wrote and revised the manuscript.

## Code availability

The code used for data processing and analysis in this study is publicly available at the GitHub repository https://github.com/JDAguirre2795/seqWORM/tree/Main

